# Emergence of mosaic plasmids harboring Tn*1546*-*ermB* element in *Staphylococcus aureus* isolates

**DOI:** 10.1101/294348

**Authors:** Tsai-Wen Wan, Yu-Tzu Lin, Wei-Chun Hung, Jui-Chang Tsai, Yu-Jung Liu, Po-Ren Hsueh, Lee-Jene Teng

## Abstract

Antimicrobial resistance in *Staphylococcus aureus* is a major problem and the acquisition of resistance genes may occur by horizontal gene transfer (HGT). The transposon, an important means of HGT, is recognized as a mobile genetic element that can integrate in plasmids, replicate and transfer to other strains. We have previously reported a novel structure of the *Enterococcus faecium*-originated Tn*1546*-*ermB* element in *S. aureus*. The emergence of the Tn*1546*-like element is an emerging problem that requires continuous monitoring. In the present study, we expand the examination of Tn*1546*-*ermB* element to *ermB*-positive methicillin-susceptible *S. aureus* (MSSA) (n = 116) and *ermB*-positive methicillin-resistant *S. aureus* (MRSA) (n = 253) during a 16-year period, from 2000 to 2015. PCR mapping showed that 10 MSSA and 10 MRSA carried the Tn*1546*-*ermB* element. The 10 MSSA belonged to three sequence types (ST), ST7 (n = 6), ST5 (n = 3), and ST59 (n = 1), and the 10 MRSA belonged to two STs, ST188 (n = 8) and ST965 (n = 2). Since only clonal complex 5 (including ST5, ST85, ST231, and ST371) MRSA, ST8 MRSA and ST5 MSSA have been previously reported to carry Tn*1546* plasmids, this is the first report describing the presence of the Tn*1546*-*ermB* element in ST7/5/59 MSSA and ST188/965 MRSA. Plasmid sequencing revealed that the Tn*1546*-*ermB* element was harbored by five different mosaic plasmids. In addition to resistance genes, some plasmids also harbored toxin genes.

## INTRODUCTION

Antimicrobial resistance in *Staphylococcus aureus* is a major problem due to its high capacity to acquire drug resistance genes by horizontal transfer (1). The problem of multidrug-resistant *S. aureus* is becoming more serious, and the available drugs to combat it are decreasing (2).

Antibiotic resistance is highly prevalent among common pathogenic bacteria in Taiwan. For example, the rate of oxacillin resistance in 33,305 *S. aureus* isolates from eight medical centers in Taiwan is approximately 60%, and the rates of resistance to erythromycin, clindamycin, gentamicin, and tetracycline are also very high (3). Erythromycin, a macrolide class antibiotic, is an old and well-established antimicrobial agent that was approved in Taiwan in 1968 (4). Due to its long history of usage in Taiwan, *S. aureus* isolates are highly resistant to erythromycin (3).

There are two major mechanisms for resistance to erythromycin. One uses methylase (encoded by *erm*) which methylates 23S rRNA, thereby altering the drug binding site, thus conferring resistance not only to macrolides (erythromycin) but also to lincosamides (clindamycin) and streptogramin B (MLS phenotype) antibiotics. The common *erm* gene in *S. aureus* is in Tn*554*-*ermA* (5, 6), Tn*551*-*ermB* (7, 8), and *ermC* (5). The second mechanism is through an efflux pump encoded by the *msrA* or *msrB*, which confers resistance to the macrolides and streptogramin B only (MS phenotype) (9).

We previously reported that the most prevalent resistance gene in erythromycin resistant blood isolates of methicillin-susceptible *S. aureus* (MSSA) collected from 2000 to 2012 in Taiwan was *ermB* (10). Although the majority (92%) of *ermB*positive MSSA isolates carried structures resembling the mobile element structure (MES) that has been reported in sequence type 59 (ST59) MRSA (8, 11), we found a unique structure, the Tn*1546*-*ermB* element (10). The significance of the Tn*1546-ermB* element is that Tn*1546* is also responsible for vancomycin resistance in *Enterococcus* spp. (12, 13). The Tn*1546*-*vanA* was mainly located on the RepA_N (pRUM/pLG1) and Inc18 plasmid families in vancomycin-resistant *E. faecium* (VREfm) (14). The plasmids RepA_N (pRUM/pLG1) were able to yield plasmid mosaics and acquire Tn*1546*-*vanA*, which was the main reason that *vanA* spread in VREfm (14).

Tn*1546* could acquire different insertion sequences (ISs), including the most frequently acquired IS*1216V*, as well as IS*1542*, IS*1251*, and IS*19* (15-18). A 92-kb plasmid of *E. faecium*, pHKK701, contains a 39-kb Tn*5506*, which carries *ermB* and IS*1216V*, and contains Tn*1546* with *vanA* (8, 19). The novel Tn*1546*-*ermB* element, Tn*1546*-carrying Tn*551*-*ermB* and IS*1216V* was recently reported in *S. aureus* by our group (10).

In this study, we examined the prevalence of the Tn*1546*-*ermB* element in MSSA and MRSA during a 16-year period, from 2000 to 2015. The presence of a Tn*1546*like element in different plasmids and lineages of *S. aureus* is an indication that *S. aureus* may have the chance to acquire Tn*1546*-*vanA* by horizontal gene transfer (HGT).

## RESULTS

### Prevalence of the Tn*1546*-*ermB* element

Of 340 erythromycin-resistant MSSA isolates collected between 2000 and 2015, 112 (33%) carried the *ermB* gene. Of 1429 erythromycin-resistant MRSA isolates collected between 2006 and 2015, 224 (16%) carried the *ermB* gene. However, only 10 MSSA and 10 MRSA carried the Tn*1546*-*ermB* element.

### Molecular epidemiology of the Tn*1546*-*ermB* element-carrying isolates

To determine the clonal relation between 10 MSSA and 10 MRSA isolates carrying the Tn*1546*-*ermB* element, we performed *spa* typing, multi-locus sequence typing (MLST) (Table 1) and pulsed-field gel electrophoresis (PFGE) (Fig. 1). Of the 10 MSSA isolates, six belonged to ST7 (*spa* type t796), three belonged to ST5 (*spa* type t002 and t242) and one isolate belonged to ST59 (*spa* type t216). Of the 10 MRSA isolates, eight belonged to ST188 (*spa* type t189 and t5529) and two isolates belonged to ST965 (*spa* type t575 and t062). The results of PFGE are shown in Fig. 1. Three ST5 MSSA isolates from 2002, 2010 and 2012 samples belonged to the same pulsotype (with 80% similarity cut-off). Two ST965 (CC5) MRSA isolates obtained from 2012 and 2014 samples were very closely related to ST5 MSSA. Six ST7 MSSA isolates obtained from 2004 to 2016 samples belonged to the same pulsotype, and the four isolates from 2012 to 2015 samples were identical. Eight ST188 MRSA isolates obtained from 2009 to 2014 samples belonged to the same pulsotype, and six of them were identical.

**Table 1.** Distribution of MLST and *spa* types in Tn*1546*-*ermB* carrying MSSA and MRSA

| Organism | Year | No. of *ermB* carried (%)<sup>a</sup> | No. of Tn1546 carried (%)<sup>b</sup> | MLST (No. of isolates) | *spa* type (No. of isolates) |
| --- | --- | --- | --- | --- | --- |
| MSSA | 2000-2015 | 112 (32.9) | 10 (8.9) | ST7 (6) | t796 (6) |
|  |  |  |  | ST5 (3) | t002 (2), t242 (1) |
|  |  |  |  | ST59 (1) | t216 (1) |
| MRSA | 2006-2015 | 224 (15.7) | 10 (4.5) | ST188 (8) | t189 (7), t5529(1) |
|  |  |  |  | ST965 (2) | t575 (1), t062 (1) |

a: Percentage of *ermB* in erythromycin-resistant isolates.
b: Rate of Tn1546 in *ermB*-positive isolates.

**FIG 1.**
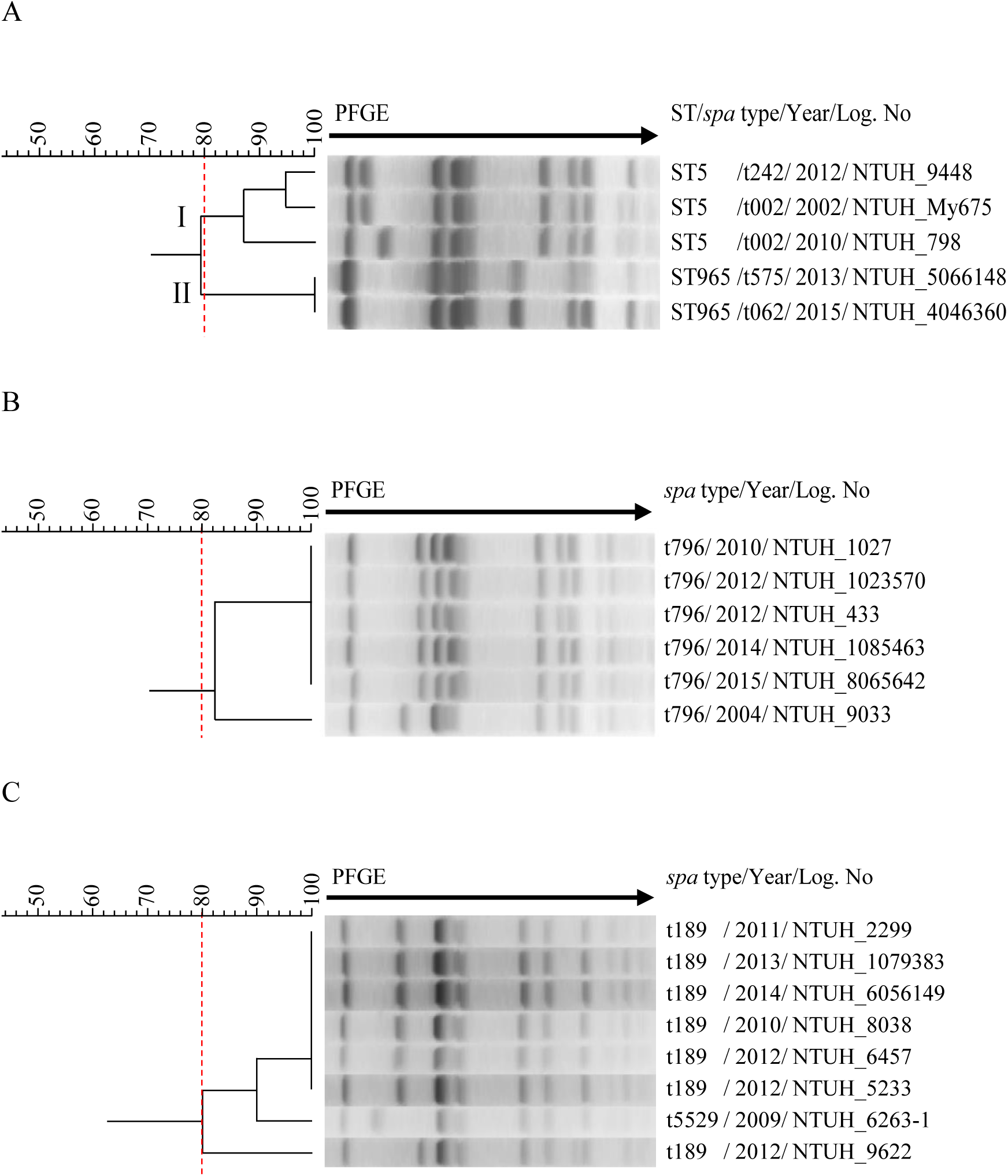
PFGE dendrogram of Tn*1546*-*ermB* carrying isolates. PFGE cluster was assigned to isolates having 80% or greater similarity from the dendrograms (A) Three ST5 MSSA isolates (pulsotype I) and two ST965 MRSA isolates (pulsotype II) (B) Six ST7 MSSA isolates (C) Eight ST188 MRSA isolates.

### Sanger sequencing of the Tn*1546*-*ermB* element and homology analysis

Representative strains of each STs (ST5 MSSA NTUH_9448, ST7 MSSA NTUH_1027, ST59 MSSA NTUH_3874, ST188 MRSA NTUH_6457 and ST965 MRSA NTUH_5066148) were chosen for sequencing to determine the sequence of Tn*1546*-*ermB* element. The size of the Tn*1546*-*ermB* element in all of the above strains is 14,567 bp. The sequences of Tn*1546*-*ermB* element in ST7 MSSA NTUH_1027 and ST965 MRSA NTUH_5066148 were 100% identical. ST5 MSSA NTUH_9448 had a nucleotide difference G1354A in *tnp* gene of Tn*551* causing G452R. ST188 MRSA NTUH_6457 had a nucleotide difference downstream of *ermB* gene. ST59 MSSA NTUH_3874 had four nucleotides difference: (1) point mutation (A299G) of *ermB* gene resulting in N100S, (2) non-coding region point mutation downstream of *ermB* gene, (3) point mutation (G765A) of *tnp* gene of Tn*551*, (4) premature nonsense mutation in *tnp* gene of Tn*551*, shortening the amino acid length from 972 to 857.

### Location of Tn*1546*-*ermB* element

To determine the location of the Tn*1546*-*ermB* element, the agarose plugs of 10 MSSA and 10 MRSA isolates were digested with S1 nuclease (Fig. 2) and then hybridized with a Dig-labeled *ermB*-specific probe prepared by PCR amplification of *ermB* using primers ermB-f and ermB-r (Table 2). Isolates of the same ST (ST5, ST7, ST188 and ST965) harbored similar size of plasmids containing *ermB* (Fig. 2).

**Table 2.**
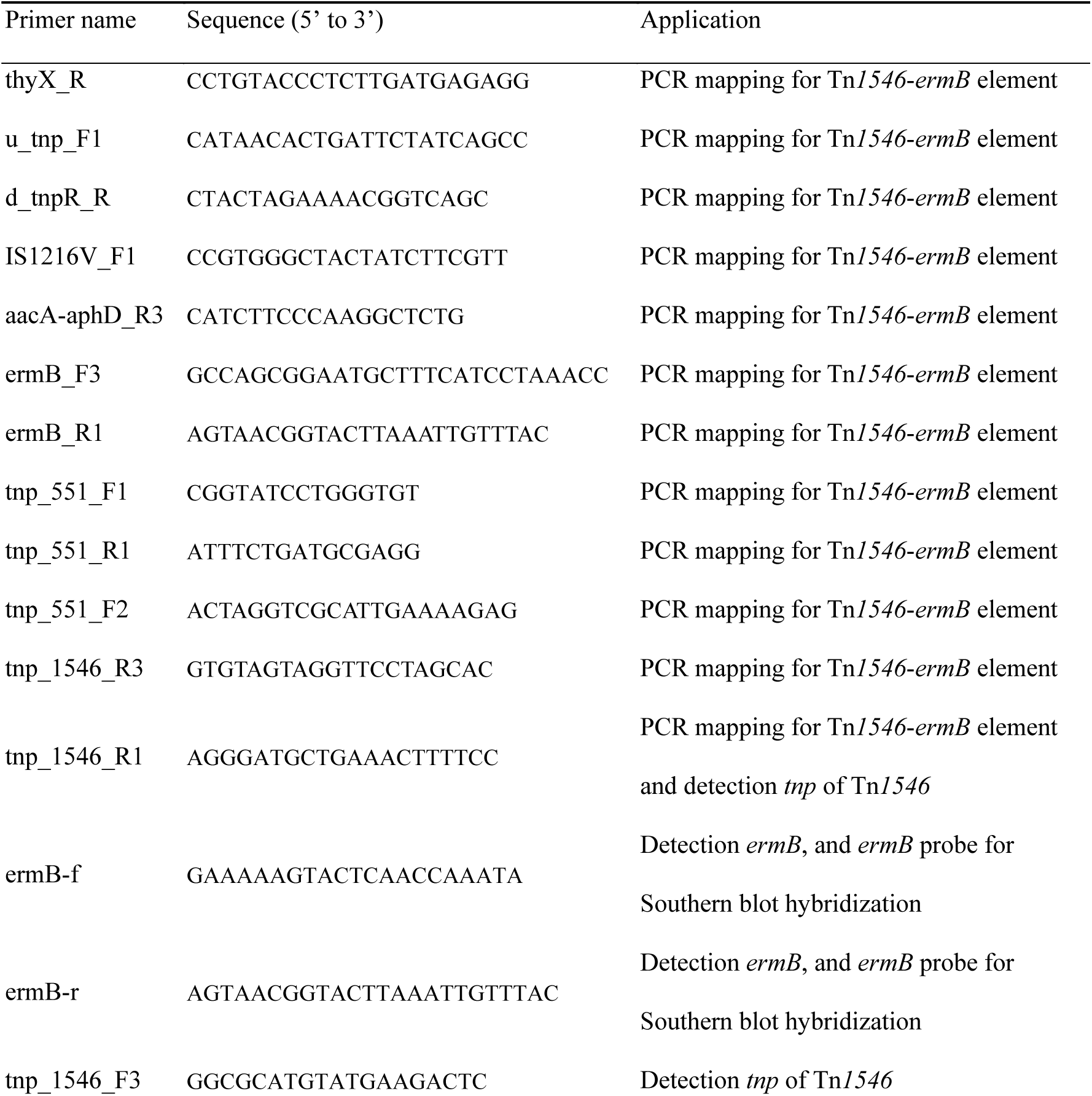
Primers used for PCR mapping of Tn*1546*-*ermB* element

**FIG 2.**
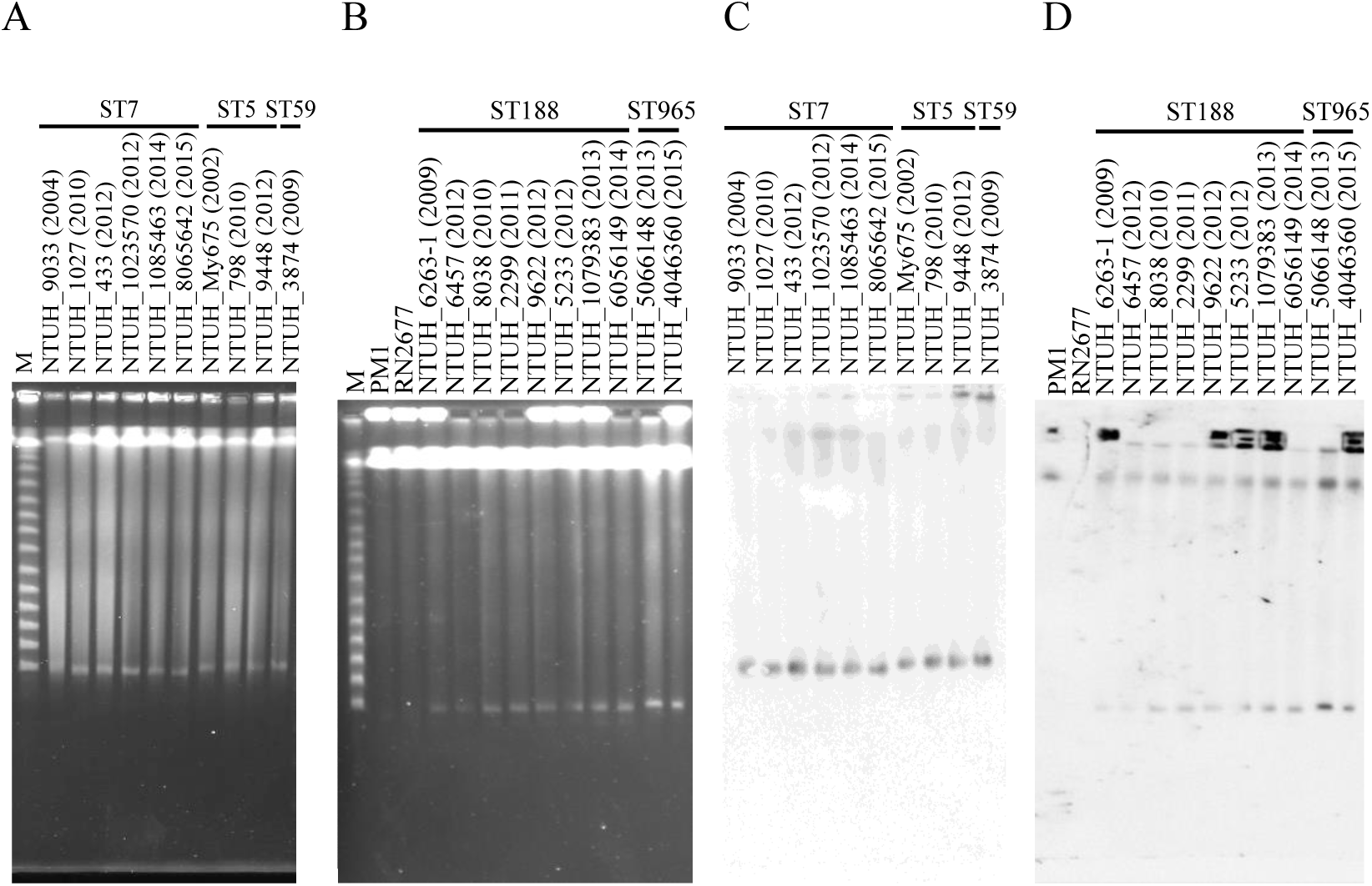
S1 nuclease PFGE and Southern blot hybridization with *ermB*. The S1 nuclease PFGE from three ST5 MSSA, six ST7 MSSA, one ST59 MSSA, eight ST188 MRSA, and two ST965 MRSA was examined. The PM1 strain harboring a 26-kb plasmid and *ermB* gene located on chromosome was used as a positive control. Strain RN2677, lacking the plasmid and *ermB* gene, was used as a negative control. The size of the M marker starts at 48.5 kb and increases 48.5 kb with each successively larger band. (A, B) The bands indicated the plasmids. The size of plasmids is estimated between 23.1 kb and 48.5 kb. (C, D) DNA was hybridized with the Dig-labelled *ermB*-specific probe and amplified by PCR using primers ermB-f and ermB-r. Positive signal of *ermB* was detected in respective bands.

### Sequence analysis of five plasmids harboring the Tn*1546*-*ermB* element

The sequences of plasmids in ST5 MSSA NTUH_9448, ST7 MSSA NTUH_1027, ST59 MSSA NTUH_3874, ST188 MRSA NTUH_6457 and ST965 MRSA NTUH_5066148 were determined. Fig. 3 presents the genetic structures of the five plasmids. The 14.5-kb Tn*1546*-*ermB* element was inserted in four different backbones. Among them, the plasmids pNTUH_1027 and pNTUH_6457 (Fig. 3A and 3B) showed mosaic structures that included Tn*1546*-*ermB* and a 20.7-kb backbone which is similar to pSaa6159 (NCBI accession no. CP002115) of ST93 MRSA. The *repA* gene and the backbone of the above two plasmids were 100% and 99.9% identical, respectively, to those in pSaa6159. The Tn*1546*-*ermB* element was inserted at the eleven o’clock position in pNTUH_1027. The Tn*1546*-*ermB* element was inserted at three o’clock position in pNTUH_6457 with the disruption of a *tnp* gene of Tn*552*. Plasmid pNTUH_5066148 (Fig. 3C) was also a mosaic plasmid and contained the Tn*1546*-*ermB* and a 24.7-kb backbone which is similar to pCA-347 (NCBI accession no. CP006045) of ST45 MRSA is essentially identical to pN315 (NCBI accession no. AP003139) of ST5 MRSA. The *repA* gene and the backbone of the pNTUH_5066148 was 100% and 99.8% identical, respectively, to those in pCA-347. The Tn*1546*-*ermB* element was inserted at the eleven o’clock position in pNTUH_5066148 with the disruption of one of the three *rep* gene, shortening the amino acid length from 286 to 271. Plasmid pNTUH_9448 (Fig. 3D) was another mosaic plasmid, in which the 14.5-kb Tn*1546*-*ermB* element was inserted at the five o’clock position and disrupted an alcohol dehydrogenase gene. The *repA* gene and the backbone of the pNTUH_9448 was 100% and 94.5% identical, respectively, to those in pWBG744.

**FIG 3.**
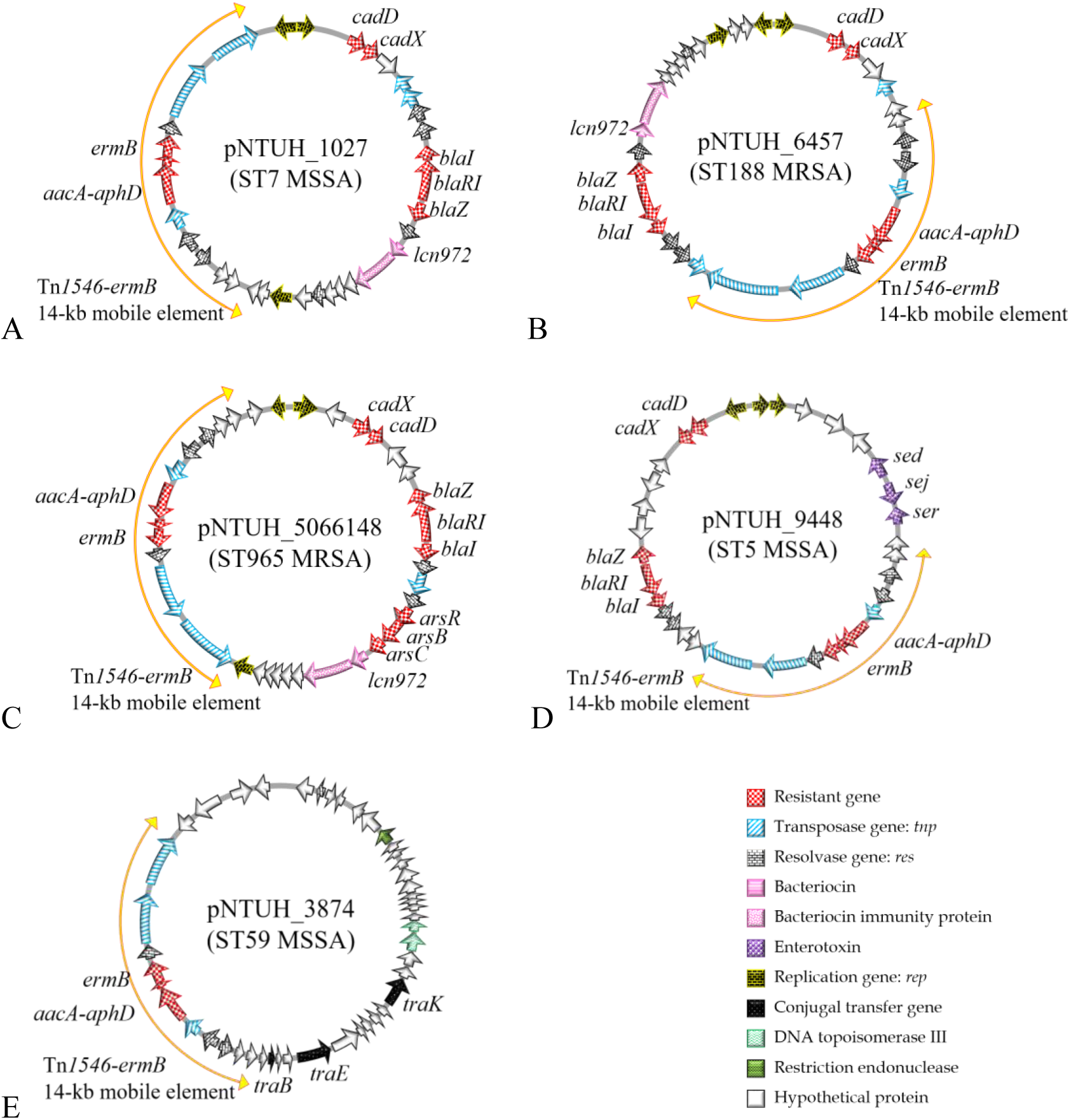
Structure of plasmids. (A) The 35.2-kb ST7 MSSA NTUH_1027 plasmid included 14.5-kb Tn*1546*-*ermB* element and 20.7-kb backbone which is similar to pSaa6159, harboring bacteriocin Lcn972 (*lcn972* gene, pink stripe). The Tn*1546*-*ermB* element was inserted at the eleven o’clock position in pNTUH_1027. (B) The 35.2-kb ST188 MRSA NTUH_6457 plasmid included 14.5-kb Tn*1546*-*ermB* element and 20.7-kb backbone which is similar to pSaa6159, harboring bacteriocin Lcn972 (*lcn972* gene, pink stripe). The Tn*1546*-*ermB* element was inserted at the three o’clock position in pNTUH_6457. (C) The 39.2-kb ST965 MRSA NTUH_5066148 plasmid contained the 14.5-kb Tn*1546*-*ermB* and a 24.7-kb backbone which is similar to pCA-347, harboring bacteriocin Lcn972 (*lcn972* gene, pink stripe). (D) The 42.5-kb ST5 MSSA NTUH_9448 plasmid included 14.5-kb Tn*1546*-*ermB* element and 27.2-kb pWBG744, harboring three enterotoxin genes *sed*, *sej*, and *ser* (purple mesh). (E) The 46.8-kb ST59 MSSA NTUH_3874 plasmid harbored three conjugal genes *traB*, *traE*, and *traK*.

Plasmid pNTUH_3874 (Fig. 3E) showed unique features. Nucleotide-nucleotide BLAST analysis found no significant matches except for the region containing the Tn*1546*-*ermB* element. Annotation revealed that Tn*1546*-*ermB* element has 11 ORFs, and the backbone of plasmid pNTUH_3874 has 35 annotated ORFs, including three conjugal transfer genes (*traB*, *traE* and truncated *traK*), one resolvase gene, one DNA topoisomerase I gene and two DNA topoisomerase III genes.

### Tn*1546*-*ermB* plasmids in other isolates

PCR mapping was used to determine whether the other Tn*1546*-*ermB*-carrying plasmids contain similar structures in each ST (Fig. S1, Table S1). Our results indicated that the plasmids in two remaining ST5 MSSA, six ST7 MSSA, seven ST188 MRSA, and one ST965 MRSA display similar plasmid structures corresponding to each ST.

### Conjugal transfer frequency of plasmids harboring the Tn*1546*-*ermB* element

To determine whether Tn*1546*-*ermB* element-carrying plasmids could be transferred *in vitro* from the clinical ST5/7/59/188/965 *S. aureus* to the laboratory strain ST8 RN2677, we performed conjugation tests. The Tn*1546*-*ermB* element-carrying plasmids could be transferred from ST5 NTUH_9448, ST7 NTUH_1027, ST59 NTUH_3874, and ST965 NTUH_5066148 to RN2677 with a frequency of 3.1×10^−10^, 10^−7^, 4.4×10^−10^, and 1.5×10^−10^ per recipient cell, respectively. Transconjugants were characterized by PCR to test for the presence of *ermB* and *tnp* of Tn*1546*, and by *spa* typing, and by erythromycin/gentamicin susceptibility testing. The results showed that the four different Tn*1546*-*ermB* plasmids could be transferred *in vitro*, and the resulting four transconjugants were resistant to erythromycin and gentamicin (Table 3).

**Table 3.** Transfer frequency of plasmids by conjugation

| Donor MSSA |  |  | Transfer frequency <sup>a</sup> | MIC of transconjugant (µg/ml) |  |
| --- | --- | --- | --- | --- | --- |
| Strain | Genotype | Plasmid size (kb) |  | Erythromycin | Gentamicin |
| NTUH_9448 | ST5/*spa* t242 | 42.5 | 3.1 X 10<sup>-10</sup> | >256 | 8 |
| NTUH_1027 | ST7/*spa* t796 | 35.2 | 10<sup>-7</sup> | >256 | 8 |
| NTUH_3874 | ST59/*spa* t216 | 46.8 | 4.4 X 10<sup>-10</sup> | >256 | 8 |
| NTUH_5066148 | ST965/*spa* t575 | 35.2 | 1.5 X 10<sup>-10</sup> | >256 | 64 |

381 <sup>a</sup> Transconjugant per donor cell.

## DISCUSSION

This study continued the work of our previous study in which the novel structure of an *E. faecium*-originated Tn*1546*-*ermB* element in MSSA was reported (10). In the present study, we examined the presence of Tn*1546*-*ermB* plasmids in more isolates of MSSA and MRSA. The overall prevalence of Tn*1546*-*ermB* plasmid in erythromycin-resistant *S. aureus* was low, and was slightly higher in MSSA than MRSA. This result is expected since most of our MRSA isolates were resistant to erythromycin and belonged to ST5/SCC*mec*II or ST239/SCC*mec*III harboring *ermA* located on Tn*554* (20).

The most frequent ST of 20 Tn*1546*-*ermB* positive isolates is ST188 SCC*mec*IVa MRSA (n = 8), followed by ST7 MSSA (n = 6). The ST188 was the common ST of MSSA in Taiwan (21). However, the ST188 in MRSA is rare in Taiwan; only one nasal carriage and one bloodstream report have referred to the ST188 MRSA (22, 23). ST188 MRSA is also rare in other countries. Recently, Hong Kong (24, 25), China (26) and Korea (27) showed the occurrence of ST188 MRSA. We examined the ST188 MRSA isolates recovered from blood from 2006 to 2012 in NTUH, and found that all the tested ST188 MRSA isolates harbored the Tn*1546*-*ermB* plasmid. The reason for this finding is unclear. PFGE analysis shows that ST188 MSSA and ST188 MRSA belong to different pulsotypes (data not shown), suggesting some level of genetic diversity.

The ST7 MSSA is one of the major sequence types of MSSA bacteremia in Taiwan (21) and is a prevalent clonotype (13/92, 14.1%) for invasive community-acquired MSSA infection (28). ST7 MSSA was the dominant MSSA carrying Tn*1546*-*ermB* element in this study.

Three ST5 MSSA and two ST965 MRSA carried the Tn*1546*-*ermB* plasmid. ST965 is a single-locus variant (SLV) of ST5. Our data shows that ST5 MSSA and ST965 harbored different plasmids. However, according to PFGE data, the pulsotypes of ST5 MSSA and ST965 MRSA were very close. Twelve cases of vancomycin-resistant *S. aureus* (VRSA) infection have been reported in the United States—all CC5 strains, each have Tn*1546*-*vanA*. The CC5 isolates appear to be very well adapted for acquiring Tn*1546* (29). In the present study ST5-CC5 MSSA and ST965-CC5 MRSA acquired Tn*1546*.

The S1 nuclease PFGE analysis showed that each Tn*1546*-*ermB* positive strain harbored only one plasmid. Isolates of the same ST carried a similar size and the same structure of plasmid. The earliest strain of this study was ST5 MSSA NTUH_My675 isolated from 2002, which indicates that the acquisition of the Tn*1546*-*ermB* element in *S. aureus* did not occur recently.

Sequencing of plasmids revealed that there were four different plasmid backbones inserted with the Tn*1546*-*ermB* element to form five mosaic plasmids. The ST7 pNTUH_1027 and ST188 pNTUH_6457 shared nearly identical plasmid backbones and best match to a 20.7-kb pSaa6159 (NCBI accession no. CP002115) derived from a dominant clone ST93 isolated in 2004 from community-associated MRSA (CA-MRSA) in Australia (30, 31). Plasmid pSaa6159 was also highly similar to the pMW2 plasmid (NCBI accession no. AP004832) from strain MW2 USA400 ST1 MRSA, which caused fatal septicemia and septic arthritis in a 16-month-old girl in North Dakota, USA, in 1998 (32). The pMW2-like plasmids are common with a wide geographical distribution (33). This is the first report that pMW2-like plasmid obtained Tn*1546*.

The best match of the plasmid backbones from ST965 NTUH_5066148 was a 24.7-kb pCA-347 (NCBI accession no. CP006045), derived from a dominant clone USA600 ST45 MRSA from a bacteremia infection in 2005 in California (34). Plasmid pCA-347 was also highly similar to the pN315 plasmid (NCBI accession no. AP003139) from strain N315 ST5 MRSA, was isolated in 1982 from the pharyngeal smear of a Japanese patient, which is prevalent in Japan and the USA (35).

The plasmids from ST7 pNTUH_1027, ST188 pNTUH_6457 and ST965 NTUH_5066148 contained the identical origin-of-transfer gene *oriT* mimic sequence of the pWBG749-family. Since the *oriT* may facilitate horizontal transmission (36–38), if this kind of plasmid contains the Tn*1546*-*ermB* element, then it may also acquire the Tn*1546*-*vanA* element. This is alarming since it raises the possibility of the occurrence of the Tn*1546*-*vanA* element in *S. aureus*.

The backbone of the plasmid in ST5 pNTUH_9448 is the 27-kb pWBG744 of ST5 MSSA. The mobilized plasmid pWBG744 belongs to the pIB485-family that has been reported in clinical and colonizing isolates of *S. aureus* (39). Plasmid pIB485 is the prototype *sed*/*sej*/*ser*-encoding plasmid of *S. aureus*. (40–42). The presence of the enterotoxin genes *sed*, *sej* and *ser* in the pIB485-family plasmids have been previously reported (33, 41).

Only one isolate of ST59 in our collection carried the Tn*1546*-*ermB* plasmid. The plasmid backbone of ST59 pNTUH_3874 is a novel plasmid, since there was no match to its nucleotide sequence in the NCBI database. This plasmid harbored three conjugal transfer genes *traB*, *traE* and truncated *traK*. Since the ST59 is the major genotype in both MSSA and MRSA in Taiwan (43), the occurrence of the Tn*1546*-*ermB* plasmid in ST59 needs more attention.

Previously the Tn*1546* element has only been found in pLW1043 (44) and pBRZ01 (45) in *S. aureus*. The *repA* gene of the four plasmids in the present study was different from that in pLW1043 (44) or pBRZ01 (45). The size of the *repA* gene is 960 bp in pLW1043, 984 bp in pBRZ01, 861 bp in pSaa6159 (backbone of pNTUH_1027 and pNTUH_6457945), 984 bp in pCA-347 (backbone of pNTUH_5066148), and 945 bp in pWBG744 (backbone of pNTUH_9948). This is the first report of the occurrence of the Tn*1546* element in new plasmids.

Although the overall prevalence of Tn*1546*-*ermB* or its plasmids in *S. aureus* isolates was low, the Tn*1546*-*ermB* element has existed from at least 2002 to the present; perhaps there are reservoirs in other species. It is known that the Tn*1546* element originated in *E. faecium*. However, the sequence of the 14.5-kb Tn*1546*-*ermB* element is more similar to that in pMCCL2 of *Macrococcus caseolyticus* (6, 10). Studies on other species may provide more information. In addition, transposon Tn*1546* is the prototype of *vanA*-carrying transposons; if *S. aureus* was able to acquire the Tn*1546*-*ermB* element, it is likely that it could obtain the Tn*1546*-*vanA* element. The transfer of vancomycin resistance to *S. aureus* has occurred *in vivo* by interspecies transfer of Tn*1546* from a co-isolate of *E. faecalis* (44). Recently, Rossi et al. reported that a conjugative plasmid carrying the Tn*1546*-*vanA* element could be transferred to other staphylococci (46). Thus, the occurrence of Tn*1546* in *S. aureus* should be monitored.

## MATERIALS AND METHODS

### Bacterial isolates

All isolates were recovered from blood. The MSSA were collected during the period between 2000 and 2015 and the MRSA were between 2006 and 2015 at the Bacteriology Laboratory, National Taiwan University Hospital, a 2,500-bed teaching hospital in northern Taiwan. Only one isolate per patient was collected in this study. *S. aureus* was identified by the Vitek2 system and *nuc* gene detection (47). Resistance to methicillin was confirmed by *mecA* PCR.

### Detection of Tn*1546*-*ermB* element structure in erythromycin-resistant isolates

The Tn*1546*-*ermB* element was initially detected by the presence of the *tnp* gene of Tn*1546* and *ermB* gene by PCR (10). The Tn*1546*-*ermB* element structure was mapped by PCR using six primer sets which are listed in Table 2, and the positions of the primers are indicated in Fig. 4. The Tn*1546*-*ermB* element structures were determined by combining the PCR mapping results and the profiles of resistance determinants.

**FIG 4.**
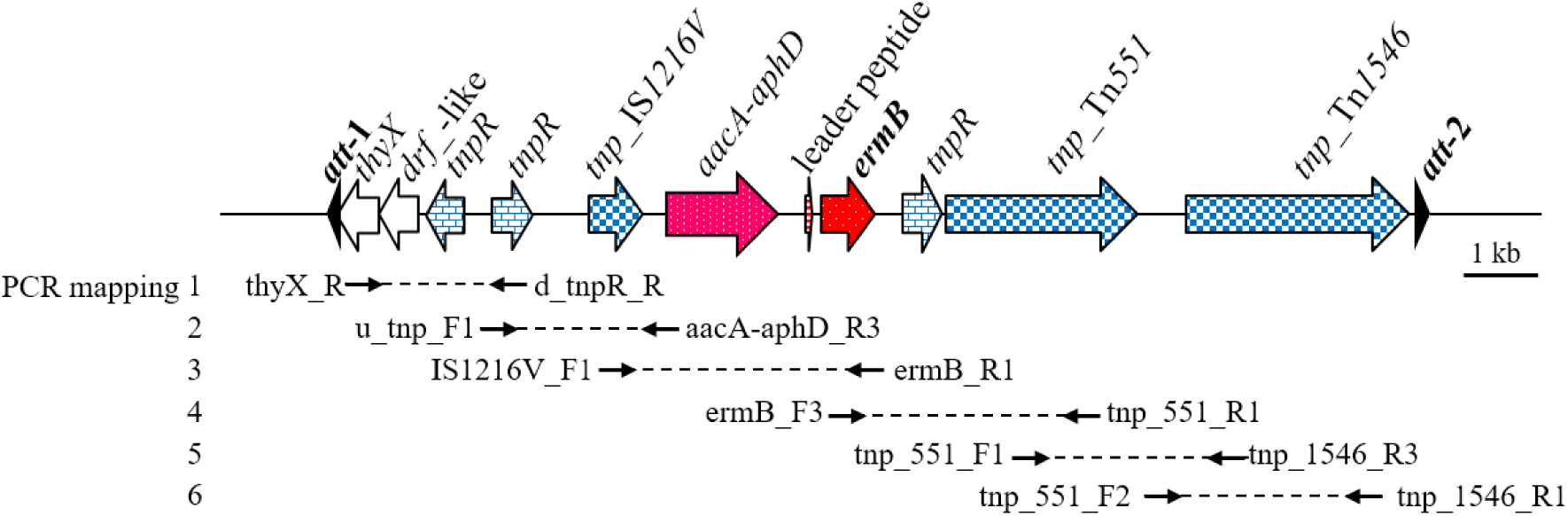
Position of PCR primers for detection of the Tn*1546*-*ermB* element structure amplicons for PCR mapping 1, 2, 3, 4, 5 and 6 are approximately 1.9 kb, 2.8 kb, 3.7 kb, 3.5 kb, 3.1 kb, and 3.2 kb, respectively.

### *spa* typing, multi-locus sequence typing (MLST), and pulsed-field gel electrophoresis (PFGE)

To determine the genetic relatedness of Tn*1546*-*ermB*-carrying isolates, *spa* typing, MLST, and PFGE were performed. The *spa* typing was performed as described previously (48). MLST was carried out to determine the sequence types (STs), which were assigned using the *S. aureus* MLST database (www.mlst.net) (49). PFGE was performed as described previously (50). The DNA in gel plugs were digested with SmaI (New England BioLabs, Ipswich, MA, USA) and then separated in a CHEF-DR III apparatus (Bio-Rad Laboratories, Hercules, CA, USA). Plugs were applied to wells in 0.8% (w/v) agarose gels (Bio-Rad). PFGE was carried out at 200 V and 12°C for 20 h, with a pulse angle of 120° and pulse times ranging from 5 to 60 s. The pulsotypes were analyzed by BioNumerics software version 4.0 (Applied Maths, Sint-Martens-Latem, Belgium).

### S1 nuclease digestion-PFGE

Detection of the presence of plasmids by S1 nuclease digestion-PFGE was performed as described previously (50). Plug slices were incubated at 37°C for 45 min with 1-10 unit of *Aspergillus oryzae* S1 nuclease (Invitrogen) in 150 μl of 50 mM NaCl, 30 mM sodium acetate (pH 4.6) and 1 mM zinc acetate. The plugs were applied to wells of 1.2% (w/v) agarose gels (Bio-Rad), and run in a CHEF-DR III apparatus (Bio-Rad Laboratories) with a pulse angle of 120° and pulse times of 45 s for 14 h and 25 s for 6 h, at 200 V in 0.5X Tris-Borate-EDTA (TBE) buffer (51). The linear form of the plasmids separated from the chromosome DNA and the size of plasmids were estimated.

### Conjugation test

To determine the transfer frequency *in vitro*, strain RN2677 was used as the recipient in the conjugation test, and mating was carried out on LB agar medium without selection (52). After 24 h, the mixed cultures were taken from the plates, suspended in brain-heart infusion (BHI) broth medium, and then plated onto MHA agar medium containing erythromycin (0.5 μg/ml) and rifampicin (80 μg/ml), at 37°C and 24 h. Confirmation of transconjugants was carried out by testing for the presence of the *ermB* gene by PCR. The transconjugants were also checked by *spa* typing (the *spa* type of RN2677 is t211) (52).

### Southern blot hybridization

DNA was electrophoresed, depurinated, denaturated, neutralized, and transferred to a Hybond-N+ nylon membrane (Amersham Pharmacia Biotech, Buckinghamshire, UK) using the Vacuum Blotting System (VacuGene^TM^ XL, Amersham Pharmacia Biotech, Buckinghamshire, UK). Hybridization with the PCR DIG Probe Synthesis Kit (Roche Diagnostics GmbH, Penzberg, Germany) was performed using a Hybridization Incubator Model 1000 (Robbins Scientific). Detection was performed with the Anti-Digoxigenin-AP and DIG Luminescent Detection Kit (Roche Diagnostics GmbH, Penzberg, Germany) and results were captured with the LAS-4000 Imaging System (FUJI FILM Life Science, Japan).

### Sequencing of plasmids

We used “Long accurate PCR *in vitro* cloning kit” (Takara Shuzo Co. Ltd., Japan) to amplify and clone the fragments of plasmids in ST5 NTUH_9448 MSSA, ST7 NTUH_1027 MSSA, ST59 NTUH_3874 MSSA, ST188 NTUH_6457 MRSA and ST965 NTUH_5066148 MRSA. The PCR products were analyzed on the ABI 3730xl DNA Analyzer (Applied Biosystems).

### Nucleotide sequence accession numbers

The nucleotide sequences of five plasmids: pNTUH_9448 in ST5 MSSA, pNTUH_1027 in ST7 MSSA, pNTUH_3874 in ST59 MSSA, pNTUH_6457 in ST188 MRSA and pNTUH_5066148 in ST965 MRSA have been deposited in the DNA Data Bank of Japan (DDBJ) database under accession numbers LC377536 to LC377540.

